# A biological spike-in enables cost-effective multi-species RNA-seq and reveals global transcriptional collapse under dark stress

**DOI:** 10.64898/2026.06.16.732598

**Authors:** Sabrina Tan, Jia Min Lee, Yan Lin Li, Chong Jun Boon, Marek Mutwil

**Affiliations:** School of Biological Sciences, Nanyang Technological University, Singapore; Department of Plant and Environmental Sciences, University of Copenhagen, Denmark; College of Horticulture, Hunan Agricultural University, Changsha 410128, China

**Keywords:** transcriptomics, RNA-seq, sample multiplexing, spike-in normalization, global transcription, dark-induced senescence, *Arabidopsis*, *Brachypodium*, *Oldenlandia*

## Abstract

RNA sequencing (RNA-seq) is the workhorse of plant functional genomics, but its per-sample cost limits the number of species, conditions and replicates that can be assayed, and its standard normalization assumes that most genes do not change and that total cellular RNA is roughly constant. Both assumptions fail when a treatment globally reprograms transcription. Here we address both problems at once. We pooled carefully weighed frozen tissue from three phylogenetically distant species, *Arabidopsis thaliana* (A), *Brachypodium distachyon* (B) and *Oldenlandia corymbosa* (O), into single RNA-seq libraries, applying a six-day dark stress to A and B while including unstressed O in every pool as an internal biological spike-in. Reads were mapped to a concatenated three-species coding-sequence index. Read assignment was essentially clean: every pure library was ≥99.8 % correctly assigned and cross-species mis-mapping was ≤0.2 %, establishing that pooling does not compromise species-level quantification. Because equal mass, not equal RNA, was pooled, the read share captured by the unchanging O reported the global RNA content of the stressed species directly: dark stress reduced total mRNA to ∼50–60 % of control in Arabidopsis and to only ∼25–37 % in Brachypodium, a magnitude difference invisible to conventional analysis. At the gene level, standard per-species analysis returned a balanced set of up- and down-regulated genes, whereas spike-in normalization revealed a response dominated by repression. Conventional pathway enrichment, measured against the bulk transcriptome, likewise failed to register the global shift. Multi-species multiplexing with a biological spike-in is therefore a cheap, quantitatively faithful strategy for stress transcriptomics.

## Introduction

RNA sequencing has become the default method for quantifying genome-wide transcript abundance, yet the cost of library preparation and sequencing still constrains experimental design, forcing trade-offs between the number of genotypes, conditions, time points and biological replicates [1,2]. Approaches that increase the information returned per library are therefore of broad value. Barcoded multiplexing of many individually-prepared libraries on one flow cell is routine, and early-pooling protocols such as BRB-seq and Lasy-Seq drive down per-sample cost by barcoding before library construction [3,4]. Pooling biological material from different species prior to extraction, however, has been little explored, largely because of the concern that reads from one species will mis-map to the genome of another and corrupt quantification. Given that library construction can in some cases amount to 50 % of the total sequencing cost, performing a single library construction for the pool rather than one per species would allow for cheaper sequencing.

A second, more fundamental limitation motivates this study. The between-sample normalization used by the dominant differential-expression tools, the median-of-ratios method of DESeq2 and the trimmed-mean-of-M-values (TMM) method of edgeR, estimates per-library size factors under the assumption that the ma_*j*_ority of genes are not differentially expressed and that total RNA per cell is roughly stable across conditions [5–8]. However, many biologically important perturbations violate this assumption. Stresses that arrest growth or shut down photosynthesis cause a genuine, genome-wide reduction in transcription, so the “average” gene is not a stable reference; under such global shifts, internal normalization can mask, or even invert, the true direction of expression change [9–13]. RNA-seq counts are, moreover, intrinsically compositional: because the read total of a library is fixed by sequencing depth, only relative abundances are measured, and a gene can appear significantly up-regulated simply because other transcripts have disappeared [14,15].

The established remedy is to anchor measurements to an external reference. Synthetic spike-in standards, most prominently the External RNA Controls Consortium (ERCC) set, are added at known amounts and allow normalization to absolute molecule numbers [16–18], and control-gene factor-analysis methods such as RUVg exploit stably-expressed genes for the same purpose [13]. Spike-ins, however, are synthetic, are introduced at the purified-RNA stage, and therefore do not experience the same extraction and library chemistry as the sample; their utility for plant studies has been demonstrated only recently [12]. Cell-number-based and total-RNA-based normalizations confirm that, once an absolute anchor is available, “hidden” global transcriptional changes become visible [9,11,19].

We reasoned that an unstressed living species, pooled into the sample as frozen tissue, could serve as a biological spike-in that experiences identical handling, extraction and sequencing while remaining transcriptionally constant because it never receives the treatment. If the species in the pool are phylogenetically distant enough that their coding sequences do not cross-map, reads can be partitioned unambiguously by species after alignment to a concatenated reference, and the read share captured by the unstressed species becomes a direct readout of how much total RNA the stressed species have lost.

To test this we used three flowering plants that span the angiosperm tree: the dicot model *Arabidopsis thaliana* [20,21], the monocot grass model *Brachypodium distachyon* [22,23], and *Oldenlandia corymbosa*, a stress-tolerant rubiaceous weed whose genome was recently sequenced [24] and which we used as the unstressed normalization control. We applied a prolonged dark stress, a treatment that halts photosynthesis, degrades chlorophyll and triggers carbon-starvation-driven transcriptional reprogramming and senescence [25–28], to Arabidopsis and Brachypodium, while the Oldenlandia tissue added to every pool was untreated. We address three questions: (i) does concatenating references and pooling species distort read mapping or species assignment; (ii) can the unstressed species quantify and correct the global change in RNA content induced by stress; and (iii) which differential-expression strategy yields biologically faithful results under this design.

## Results

### A multi-species spike-in design for Arabidopsis, Brachypodium and Oldenlandia

To test whether plant material from several species can be co-sequenced and one species used as a normalization control, we grew the three species and exposed *Arabidopsis* (30 days after germination, DAG) and *Brachypodium* (15 DAG) to six days of continuous darkness; *Oldenlandia* (∼30 DAG) was never dark-treated. As expected for dark stress, treated A and B plants became chlorotic (Figure 1A), and pigment quantification showed significant reductions in chlorophyll *a* and chlorophyll *b* in both species, and in carotenoids in Arabidopsis; the change in Brachypodium carotenoids was not significant (Figure 1B; Table S1) [25,26]. We then combined carefully weighed 100 mg masses of frozen powder from the three species in three biological replicates per condition (Figure 1C). Each pool was sub_*j*_ected to a single RNA extraction and Illumina sequencing, and reads were mapped with HISAT2 [29] to the concatenated coding sequences (CDS) of all three species; mapped reads were counted per transcript with SAMtools idxstats [30,31].

**Figure 1.**
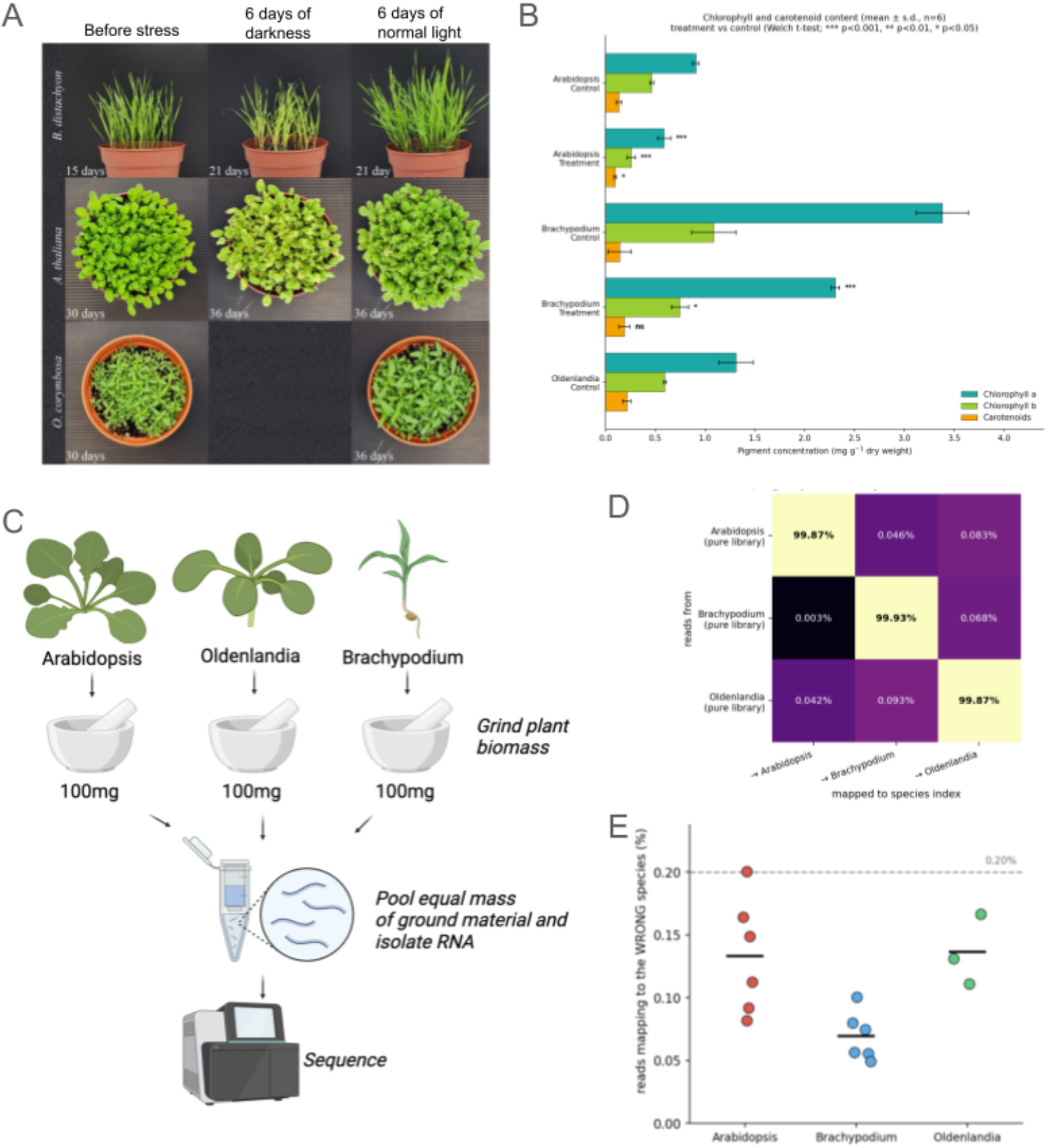
Multi-species pooling design and multiplexing fidelity. (A) *B. distachyon, A. thaliana* and *O. corymbosa* before treatment, after six days of dark stress, and untreated controls. (B) Chlorophyll *a*, chlorophyll *b* and carotenoid content (mg g^−1^ dry weight) in control and treated plants (mean ± s.d., *n* = 6; Welch *t*-test, *** p < 0.001, ** p < 0.01, * p < 0.05; Table S1). (C) Pooling scheme: equal frozen masses (100 mg per species) ground, pooled, RNA-extracted once and sequenced, then mapped to a concatenated three-species coding-sequence index. Figure panel generated with Biorender. (D) Confusion matrix of read assignment: each pure single-species library mapped to the concatenated index; rows = source species, columns = index mapped to, values = % of reads (diagonal ≥ 99.87 %). (E) Cross-species mis-mapping per library (off-species read share, %). Each circle indicates on RNAseq sample.

### Pooling and reference concatenation do not distort species-level quantification

Mapping each pure single-species library to the concatenated three-species index showed that reads are partitioned almost perfectly by species: ≥99.8 % of reads from every library mapped to the correct species’ coding sequences (Figure 1D), and the worst-case cross-species assignment for any pure sample was 0.20 %, with most libraries below 0.13 % (Figure 1E). Mis-mapping was lowest between the most divergent pair: only 0.003 % of *Brachypodium* reads mapped to *Arabidopsis*, confirming that phylogenetic distance drives the clean separation. A splice-aware aligner with a single-best-hit policy therefore partitions reads cleanly across a dicot, a grass and a rubiaceous eudicot, and multiplexing mixed frozen tissue is quantitatively safe for distantly related species.

### The biological spike-in turns compositional RNA-seq into a measure of relative transcriptome size

Because equal frozen mass rather than equal RNA was pooled, the proportion of reads captured by each species reflects its mRNA content per unit tissue (Figure 2A; mapped reads in Table S2). Averaged across replicates, the *Oldenlandia* read share rose from 36 % in the control pools to 63 % under dark stress (composition_pct in Table S2), while *Brachypodium* fell from ∼49 % to ∼22 % and *Arabidopsis* changed little (∼15 %) (Figure 2B). Since an identical, untreated mass of Oldenlandia was present in every pool, its share cannot have risen because Oldenlandia changed; it rose because the stressed transcriptomes of Arabidopsis and Brachypodium shrank around it (per_sample_ratios in Table S2). Anchoring each pool to the constant Oldenlandia therefore converts the compositional readout into an estimate of per-species mRNA mass (Figure 2C): the combined transcriptome of the stressed pair fell to roughly one third of control, with Arabidopsis retaining ∼50–60 % of its control mRNA and Brachypodium only ∼25–37 % (Table S2).

**Figure 2.**
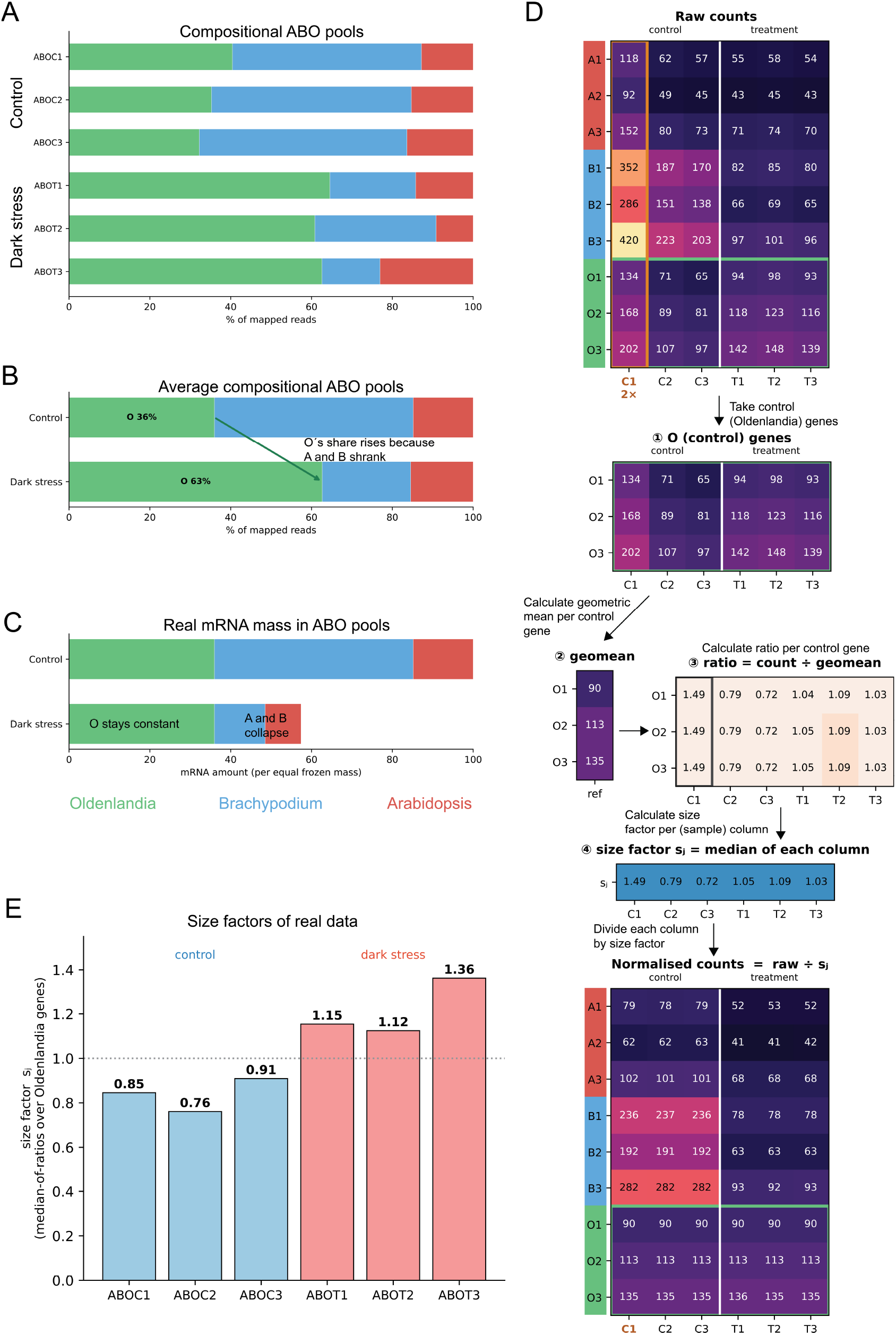
A biological spike-in turns compositional RNA-seq into a measure of transcriptome size. (A) Species composition (% of mapped reads) of the six three-species (ABO) pools. ABOC and ABOT are Control and Treated (Dark Stress), respectively. (B) Average composition, control versus dark stress. (C) The same data re-expressed as mRNA mass relative to the spike-in, by anchoring to the constant *Oldenlandia*. (D) Control-gene normalization mechanism: per-sample size factors *s*_*j*_ are estimated by median-of-ratios over the *Oldenlandia* genes only (raw counts → per-gene geometric-mean reference → ratios → median per sample) and applied to every gene, leveling the spike-in. (E) Empirical control-gene size factors for the six ABO pools (controls < 1 < dark stress).

Thus, the total mRNA pools of Arabidopsis and Brachypodium decreased dramatically under dark stress, indicating a genome-wide downregulation of gene expression.

### Control-gene size factors normalize to the constant spike-in

The spike-in revealed a transcriptome collapse in the dark-stressed plants, but to use this information for differential gene expression (DGE) analysis, we need to modify the typical normalization pipeline. Standard DGE analyses assume that most genes do not change, but it is clear that this assumption is not valid in dark-stressed plants (Figure 2C). However, we can use genes whose expression did not change, the Oldenlandia spike-in, as a normalization reference. Figure 2D works the calculation through on a small illustrative count table.

Beginning with the raw counts for all three species (Figure 2D, top), we keep only the Oldenlandia control genes (O1-O3, step ①), because only they are invariant. For each Oldenlandia gene we calculate its geometric mean across the six samples (step ②). Dividing each gene count by the geometric mean gives a ratio (step ③), which shows that the different control genes give almost the same ratios in each sample (close to 1.49 in the first sample, close to 0.79 in the second), even though those genes are expressed at different absolute levels. This is because anything that inflates or deflates a whole library (whether deeper sequencing or the larger read share left behind as the stressed species shrink) multiplies every gene in that sample by the same amount, and it is this common factor that survives when each gene is divided by its own mean.

The size factor *s*_*j*_ for a sample is therefore the median of its column of ratios (step ④); using the median rather than the mean means that one unusual control gene cannot distort it. Dividing every gene, in all three species, by its sample’s size factor (Figure 2D, bottom) brings the Oldenlandia counts to the same level in every sample. The samples are now normalized by the spike-in, so the Arabidopsis and Brachypodium genes are read against a fixed reference rather than against one another.

Two features make this normalization robust. Because each control gene is compared only with itself, the control genes need not be expressed at similar levels, only to be unchanged by the treatment. And because a more deeply sequenced library scales all of its counts up together, the same step also removes ordinary differences in sequencing depth (in the illustration, the first control sample was sequenced twice as deeply and receives a size factor roughly twice that of the other controls, which cancels the extra depth exactly).

On the experimental data, the Oldenlandia size factors separate cleanly by condition, 0.76–0.91 for the three control pools and 1.12–1.36 for the three dark-stress pools (Figure 2E). The dark-stress libraries therefore carry size factors about 1.45 times larger than the controls.

### Spike-in normalization rescales differential expression and exposes a directional artifact

The transcriptomic collapse observed in Arabidopsis and Brachypodium indicates a global downregulation of gene expression (Figure 2A,C), while the size factors capture how much each transcriptome shrank relative to the constant Oldenlandia. DESeq2 is a popular method to calculate differential gene expression (DGE), but when used with default settings it calculates its own size factors, which override the size factors found by our spike-in scaling.

To compare the behavior of DESeq2 with or without our spike-in scaling, we (i) let DESeq2 use its internal scaling (naïve analysis) and (ii) supplied it with our size factors (spike-in scaling; Table S3). As expected from a compositional analysis, the naïve analysis produced a roughly equal number of up- and down-regulated genes, with a higher number of down-regulated genes for Arabidopsis and Brachypodium (Figure 3A). Conversely, spike-in normalization changed both the magnitude and the direction of the differential-expression result (Figure 3A,B). Relative to the naïve analysis (each species normalized to its own library, i.e. no scaling), control-gene normalization shifted every gene’s log_2_ fold-change down by a near-rigid, species-specific amount, about −1.05 in Arabidopsis and −2.23 in Brachypodium. This lowered the mean log_2_ fold-change from −0.46 to −1.50 in Arabidopsis and from −0.75 to −2.98 in Brachypodium (Figure 3B). Because this rigid shift slides genes across the |log_2_FC| > 1 threshold, apparent up-regulation collapsed while down-regulation grew: up-regulated DEGs fell from 3,828 to 1,577 (Arabidopsis) and from 4,315 to 888 (Brachypodium), and down-regulated DEGs rose from 5,338 to 9,013 and from 7,189 to 19,735, respectively (Figure 3A; Table S3). Given that the transcriptional collapse has been observed in Arabidopsis and Brachypodium, much of the up-regulation seen in a naïve analysis is thus a compositional illusion that the biological spike-in removes.

**Figure 3.**
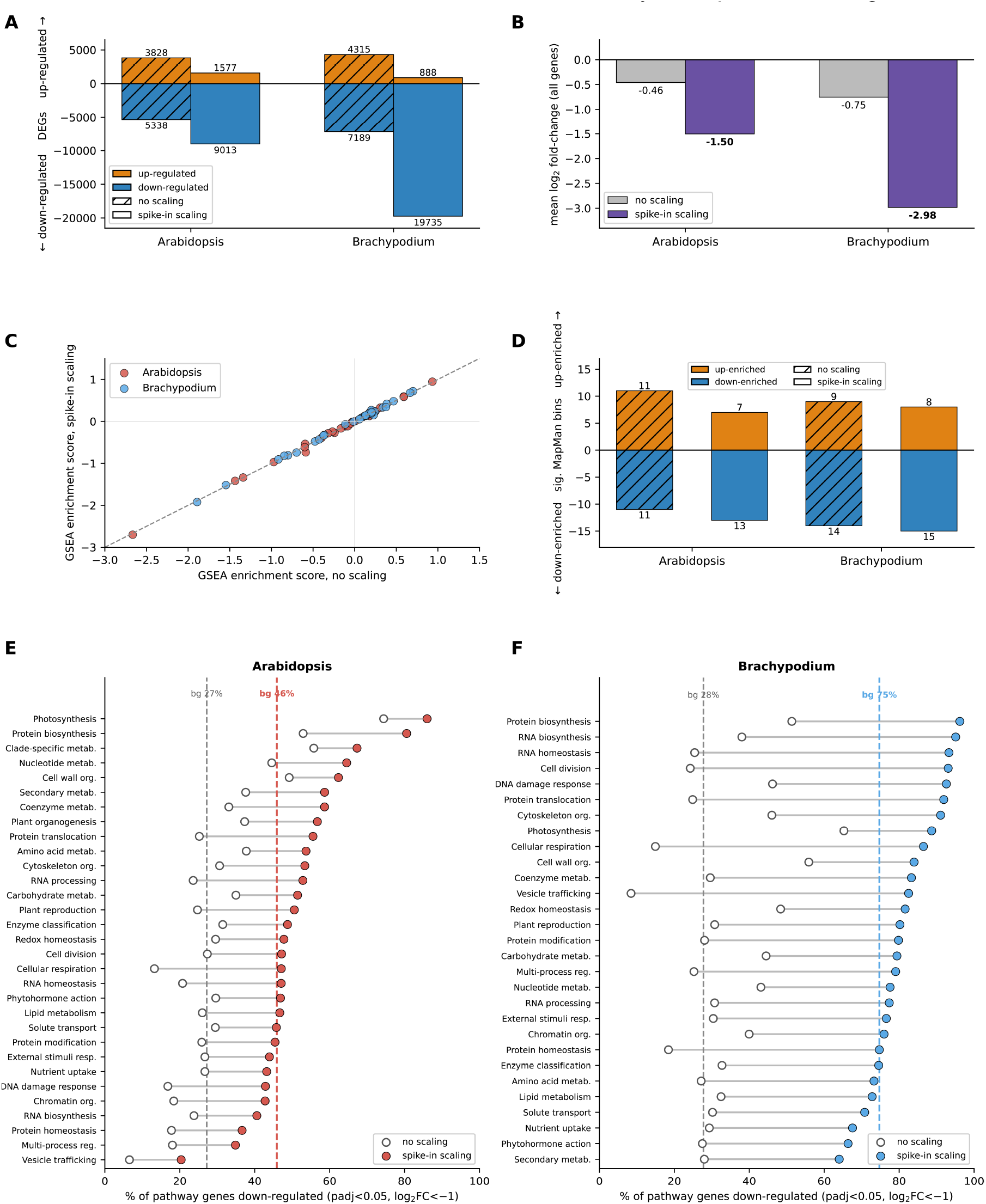
Spike-in normalization rescales differential expression; rank enrichment is invariant while threshold enrichment tracks the global shift. (A) DEGs (ad_*j*_usted p < 0.05, |log_2_FC| > 1; up/down) for *Arabidopsis* and *Brachypodium* analysed alone, without versus with spike-in scaling. (B) Mean log_2_FC across all genes per species, no scaling versus spike-in scaling. Rank-based GSEA enrichment scores per MapMan bin, no scaling versus spike-in scaling. Number of significantly over-represented MapMan bins (threshold ORA), up- and down-enriched, no scaling versus spike-in scaling. (E,F) Per-pathway down-content (% of a pathway’s genes called down-regulated), no scaling versus spike-in scaling, with the genome-wide background (dashed); pathways shift down in lockstep with the background, for *Arabidopsis* (E) and *Brachypodium* (F).

### Global transcriptome change confounds standard pathway enrichment

Differential gene expression analysis is typically followed up by a pathway enrichment analysis, to identify biological pathways that are significantly up- or down-regulated due to the perturbation. To investigate how our scaling approach affects the identified pathways, we use two standard but fundamentally different enrichment tests.

The first is a rank-based test (Gene-Set Enrichment Analysis [35], implemented here as a Mann–Whitney test). It uses no cutoff: it ranks every gene by its fold-change and asks, for each MapMan pathway, whether that pathway’s genes sit preferentially toward the up-regulated or the down-regulated end of the ranking. It therefore depends only on the relative order of genes, not on the size of their fold-changes, and we expect this method not to be affected by our correction.

The second is a threshold-based over-representation test (a hypergeometric test). It works in two stages: it first selects a list of differentially expressed genes using fixed cutoffs (typically ad_*j*_usted p < 0.05 and |log_2_FC| > 1), then determines whether a pathway contains more of those genes than expected by chance, given how many genes passed the cutoff across the whole genome. It therefore depends both on the cutoffs and on the genome-wide rate of differential expression. Because our correction down-shifted the log_2_FC dramatically for both species, we expected this method to be affected by our correction, and to observe a higher number of significantly down-regulated pathways.

The rank-based GSEA over top-level MapMan bins [32,33] identified Photosynthesis as the most strongly down-enriched category (Table S4 for Arabidopsis, enrichment score ≈ −2.7; Table S5 for Brachypodium), together with cell-wall organisation, protein biosynthesis, cell division and cytoskeleton organisation, while vesicle trafficking, protein homeostasis and multi-process regulation were the leading up-enriched categories, consistent with the autophagy- and proteostasis-associated remodelling of dark-induced senescence [25–27]. Because spike-in scaling is a near-constant shift that preserves gene ranks, rank-based enrichment scores were essentially identical with and without scaling: every MapMan bin fell on the line of equality (Figure 3C). GSEA therefore reports which pathways deviate from the bulk but, like the bulk normalization itself, cannot register the genome-wide downregulation.

Threshold-based over-representation behaved differently (Figure 3D). Up-enriched bins shrank under scaling (Arabidopsis 11 → 7, Table S4; Brachypodium 9 → 8, Table S5). The number of *significantly* down-enriched bins, however, barely changed (Arabidopsis 11 → 13; Brachypodium 14 → 15) even though down-regulated DEGs roughly tripled. This is because over-representation is measured relative to the genome-wide background, and that background shifted in step: under scaling, the fraction of all genes called down-regulated rose from 28 % to 75 % in Brachypodium and from 27 % to 46 % in Arabidopsis. Every pathway’s down-content rose in lockstep with the background (Figure 3E,F), so few individual bins exceeded it. Over-representation against a whole-genome background is thus the pathway-level analogue of the same compositional trap that confounds gene-level normalization: it silently absorbs a global shift, and is therefore blind to the global downregulation.

Both enrichment tests share a deeper limitation that this experiment exposes. Each asks whether a pathway behaves differently from the rest of the transcriptome, so when the entire transcriptome collapses, the reference against which they measure has itself collapsed and neither reports the global downregulation. The genome-wide repression is therefore invisible to both standard tests and is recovered only by absolute, spike-in-anchored metrics, such as the per-pathway down-content in Figure 3E,F, where nearly every pathway shifts down in lockstep with the background.

## Discussion

We have shown that frozen tissue from phylogenetically distant plant species can be pooled, sequenced as a single library, and confidently de-multiplexed in silico, and that including an unstressed species in the pool turns ordinary, compositional RNA-seq into a measurement of transcriptome size. Three findings support the approach. First, cross-species mis-mapping is negligible (Figure 1D,E), so pooling is quantitatively safe for species as divergent as a dicot, a grass and a rubiaceous eudicot. Second, because equal mass rather than equal RNA was combined, the read share of the constant Oldenlandia spike-in directly reports the global RNA content of the stressed species. This revealed that dark stress dramatically shrank the transcriptomes of Brachypodium and Arabidopsis (Figure 2), a difference that within-sample normalization erases. Third, the unstressed species doubles as a negative control that exposes, and lets us correct, the directional artifact by which a shrinking transcriptome inflates apparent up-regulation (Figure 3).

The methodological message generalizes beyond this experiment. RNA-seq measures composition, not absolute abundance, and the default size-factor estimators are confounded whenever the bulk transcriptome changes [9,11,14,15]. Our control-species design serves the same purpose as ERCC spike-ins and control-gene methods such as RUVg, anchoring the data to an unchanging reference, but differs in mechanism: the reference is a whole unstressed species pooled as tissue, so it experiences the full extraction and library workflow rather than being added to purified RNA (unlike ERCC) and needs no a-priori set of stable control genes (unlike RUVg), while doubling as the multiplexing partner that lowers cost. It extends the recent demonstration that controlling for total RNA changes which plant DEGs are called [12].

Biologically, the spike-in turns the experiment into a measurement of how much global transcription each species changes. The stronger collapse in Brachypodium is consistent with its younger developmental stage (15 versus 30 DAG) and its identity as a young grass seedling with limited stored carbon, both of which would heighten sensitivity to dark-induced carbon starvation [25–28]. That the down-regulated functional categories (photosynthesis, cell wall, translation, cell division) are conserved across a monocot and a dicot, while the magnitude of the response differs markedly, illustrates the value of being able to compare the size and not merely the shape of two transcriptional responses in one experiment.

Several limitations should be borne in mind. We measured mRNA per unit frozen mass relative to the spike-in, which conflates per-cell transcription with stress-induced changes in tissue water content and cell density, and which cannot distinguish reduced transcription from increased RNA turnover. Species and developmental stage are also confounded (30 versus 15 DAG), so the stronger Brachypodium collapse cannot be attributed to species identity alone. The clean de-multiplexing we observe depends on phylogenetic distance, and more closely related species would likely result in a higher rate of cross-mapping. Finally, replicate-level variation in the species ratios was appreciable, reflecting the precision of weighing frozen powder; the spike-in corrects sequencing depth but not pooling-mass error, so accurate weighing and adequate replication remain essential.

In sum, multi-species multiplexing with a biological spike-in offers a route to easier, cheaper and quantitatively honest stress transcriptomics: it lowers cost by sequencing several species in one library, it recovers per-species differential expression faithfully, and, uniquely among routine designs, it measures the global change in transcriptome size that conventional analysis discards.

## Materials and Methods

### Plant material and dark-stress treatment

*Arabidopsis thaliana* (Col-0), *Brachypodium distachyon* (Bd21-3) and *Oldenlandia corymbosa* were grown in a Percival phytotron under a 12 h light / 12 h dark photoperiod (100 µE) at 24 °C. Arabidopsis (30 DAG) and Brachypodium (15 DAG) were transferred to continuous darkness for six days (dark-stress/treatment); matched plants kept under the normal photoperiod served as no-stress controls. Oldenlandia (∼30 DAG) was grown only under control conditions.

### Pigment quantification

Chlorophyll and carotenoids were extracted from dried tissue in 95 % ethanol, and absorbance was measured at 470, 649, 655 and 665 nm. Chlorophyll *a*, chlorophyll *b* and total carotenoid concentrations were calculated using standard equations and normalized to dry weight, with six biological replicates per species and condition (Table S1). For each pigment, treated plants were compared with same-species controls by a two-sided Welch’s two-sample *t*-test (unequal variance; SciPy [38]), with no correction for multiple comparisons; significance is indicated in Figure 1B as * p < 0.05, ** p < 0.01 and *** p < 0.001.

### Multiplexing, RNA extraction and sequencing

Frozen tissue was ground to powder and equal masses of 100 mg per species were combined into one pool. Total RNA was extracted once per pool with the Sigma Spectrum Plant Total RNA Kit, poly-A enriched, and sequenced on an Illumina NovaSeq 6000 instrument in paired-end 150 bp mode by Novogene. Each control and treatment condition was performed in triplicate.

### Read mapping and counting

Reference coding sequences (primary transcript only) were obtained from Phytozome [36]: *Arabidopsis* (Athaliana_167/Araport11) [21], *Brachypodium* (Bd21-3 v1) [23] and *Oldenlandia corymbosa* (OLC1) [24]. A concatenated three-species CDS index and single-species indexes were built with hisat2-build. Reads were aligned with HISAT2 v2.2.1 [29] using -k 1 to retain only the single best alignment per read; alignments were sorted and per-transcript mapped/unmapped read counts obtained with SAMtools v1 idxstats [30,31]. Reads were assigned to species by transcript-ID prefix (At*, Bd*, OLC*). Cross-mapping was quantified as the fraction of reads from a single-species library assigned to another species on the concatenated index (Figure 1D,E).

### Spike-in (ratio) normalization and transcriptome-size estimation

For each species the species-to-Oldenlandia read ratio was computed per condition; the treatment/control ratio of these values gives the fraction of control mRNA retained under stress (Figure 2C; Table S2). Because the same untreated Oldenlandia mass is present in every pool, this ratio is a depth-independent estimate of relative transcriptome size.

### Differential expression

Counts were analysed with DESeq2 [5] (via pydeseq2 [37]); a gene was called a significant DEG at ad_*j*_usted *p* (Ben_*j*_amini–Hochberg [39]) < 0.05 and |log_2_FC| > 1. Two normalizations were compared: (i) default (median-of-ratios over all genes); and (ii) control-gene normalization, in which per-sample size factors were estimated by the median-of-ratios method restricted to the Oldenlandia genes (DESeq2 control_genes; equivalent in spirit to RUVg [13]) and fixed in the model.

### Formal definition of the normalization

Let *k*_i*j*_ be the number of reads assigned to transcript *i* in sample *j* (SAMtools idxstats, mapped column), and let *A*, ℬ, *O* denote the *Arabidopsis, Brachypodium* and *Oldenlandia* gene sets.

#### Cross-mapping

For a single-species library of species *x* aligned to the concatenated index, the cross-mapping rate is

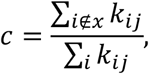

which was below 0.2 % for every pure sample.

#### Control-gene (spike-in) size factors

Applying the median-of-ratios principle restricted to the unstressed control genes *O*, define the per-gene geometric-mean reference across the *n* samples and the size factor of sample *j*:

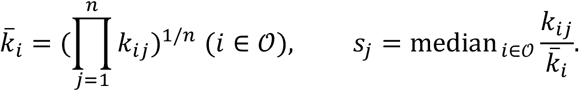

Counts are normalized as 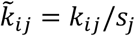 . Because *s*_*j*_ is estimated from *O* alone, 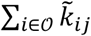 is approximately constant across samples (the spike-in is pinned), so that changes in *A* and ℬ are measured relative to a fixed control. In DESeq2 the *s*_*j*_ are supplied as fixed size factors (the control_genes option).

#### Transcriptome-size (ratio) estimate

The read share of a stressed species *s* ∈ {*A*, ℬ} relative to the constant control is *ρ*, = ∑_i∈s,_ *k*_i_ / ∑_i∈*O*_ *k*_i_ (summed over a condition’s replicates), and the fraction of control-level mRNA retained under stress is

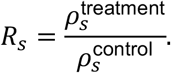

#### Significance

A gene was a DEG at Ben_*j*_amini–Hochberg *p*_ad*j*_ < 0.05 and |log_2_FC| > 1, where for *m* ordered tests 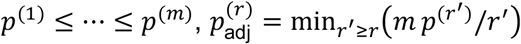.

### Functional enrichment

Coding sequences were annotated to MapMan bins with Mercator4 [33,34] and genes assigned to their top-level bin (bins with ≥10 detected genes retained). Two complementary tests were applied per species and per normalization (Figure 3C–F). For rank-based enrichment (GSEA-type [35]), the log_2_ fold-changes of in-bin genes were compared with the background by a two-sided Mann–Whitney *U* test, with sign taken from the median difference and Ben_*j*_amini–Hochberg correction across bins; this test depends only on gene ranks and is therefore invariant to the constant log_2_FC shift introduced by normalization. For threshold-based over-representation (ORA), the probability of ≥ *k* DEGs in a bin of *K* annotated genes given *n* DEGs among *N* background genes is the hypergeometric tail 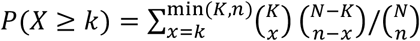, computed separately for up- and down-regulated DEGs and BH-corrected. Per-pathway down-content (fraction of a bin’s genes called down-regulated) was compared with the genome-wide background fraction to interpret the ORA result under a globally shifting transcriptome (Figure 3E,F).

## Code and data availability

Raw reads are deposited under E-MTAB accession. Analysis scripts and intermediate results are available at https://github.com/mutwil/multiplex-rnaseq-spikein.

## Supplemental Tables

**Table S1. Chlorophyll and carotenoid content.** Photosynthetic pigments were extracted from dried tissue in 95% ethanol and quantified by absorbance as chlorophyll *a*, chlorophyll *b* and total carotenoids (mg per g dry weight), with six biological replicates per species and condition. The sheet contains three tables: measurements — raw per-replicate values for all five groups (*Arabidopsis* control and treatment, *Brachypodium* control and treatment, *Oldenlandia* control), one row per replicate, with columns for species, condition, replicate number, and the three pigment concentrations. summary_mean_sd — mean and standard deviation (n = 6) of each pigment for every species and condition. welch_tests — two-sided Welch’s two-sample *t*-test (unequal variance) of treatment versus control for each species and pigment, reporting the control and treatment means, the *p*-value, and a significance code (*** p < 0.001, ** p < 0.01, * p < 0.05, ns not significant); no correction for multiple comparisons was applied. Chlorophyll *a* and *b* decreased significantly in both species and carotenoids in *Arabidopsis*, whereas *Brachypodium* carotenoids did not change significantly (p = 0.40).

**Table S2. Mapped-read counts and Oldenlandia-normalized transcriptome-size estimates for the three-species (ABO) pools.** Reads from each pool were mapped to the concatenated three-species coding-sequence index and assigned to a species by transcript-ID prefix, for the six three-species pools (three control, three dark stress). The workbook contains four tables: **mapped_reads**: number of reads assigned to *Arabidopsis, Brachypodium* and *Oldenlandia*, and the total mapped, for each pool. **composition_pct**: the same data expressed as the percentage of mapped reads contributed by each species in each pool, with the per-condition means. **per_sample_ratios**: for each pool, the reads from each stressed species expressed relative to the *Oldenlandia* reads in the same pool. Because an equal, untreated mass of *Oldenlandia* was pooled into every library, this is a sequencing-depth-independent measure of each stressed species’ mRNA content per unit tissue. **transcriptome_size**: the estimated fraction of control mRNA retained under dark stress for each stressed species, computed as the ratio of its mean read share relative to *Oldenlandia* under dark stress to the same quantity under control. Values are given with a 95% bootstrap confidence interval (resampling the three replicates per condition, 10^5^ iterations) and the treatment-group coefficient of variation. *Arabidopsis* retained 0.59 of its control mRNA (95% CI 0.36 to 0.90; CV 45%) and *Brachypodium* 0.25 (95% CI 0.17 to 0.36; CV 38%).

**Table S3. Differentially expressed gene (DEG) counts and fold-change by species, normalization and direction.** Each species was demultiplexed and analysed on its own counts with DESeq2 (pydeseq2). A gene was scored as a DEG at Ben_*j*_amini-Hochberg ad_*j*_usted *p* < 0.05 and |log_2_FC| > 1 for the dark-stress versus control contrast. Two normalizations are compared: “no scaling”, in which each species is normalized to its own library by the default median-of-ratios method (the standard analysis a researcher would run after demultiplexing), and “spike-in scaling”, in which per-sample size factors estimated from the *Oldenlandia* control genes are fixed in the model. For each species and normalization the table gives the number of up- and down-regulated DEGs, the total, and the mean and median log_2_ fold-change across all tested genes.

**Table S4. MapMan pathway enrichment in *Arabidopsis* under dark stress, by enrichment method, normalization and direction.** Each row is a top-level MapMan bin (pathways with at least 10 detected genes; *n* genes = number of detected genes in the bin), ordered from most down-enriched to most up-enriched by the spike-in GSEA score. Two complementary enrichment tests are reported, each under both normalizations: **default** (the species normalized to its own library by median-of-ratios) and **spike-in** (per-sample size factors fixed from the *Oldenlandia* control genes). **GSEA ES (default / spike-in)** — rank-based enrichment score, the median log_2_ fold-change of the bin’s genes minus that of the rest of the transcriptome (background), from a two-sided Mann-Whitney test on gene-level log_2_ fold-changes. Negative values indicate a pathway shifted toward down-regulation relative to the background, positive toward up-regulation. **GSEA FDR (default / spike-in)** — Ben_*j*_amini-Hochberg ad_*j*_usted *p*-value for that Mann-Whitney test, corrected across all bins. **ORA up FDR / ORA down FDR (default / spike-in)** — hypergeometric over-representation FDR for up-regulated and, separately, down-regulated differentially expressed genes (ad_*j*_usted *p* < 0.05 and |log_2_FC| > 1) within the bin, tested against the genome-wide background of DEGs and Ben_*j*_amini-Hochberg corrected. Shaded cells indicate FDR < 0.05.

**Table S5. MapMan pathway enrichment in *Brachypodium* under dark stress, by enrichment method, normalization and direction.** Columns as in Table S4.

## Notes

### Competing Interest Statement

The authors have declared no competing interest.

